# Synergically enhanced anti-tumor immunity of *in vivo* CAR by circRNA vaccine boosting

**DOI:** 10.1101/2024.07.05.600312

**Authors:** Yanyan Wang, Liangru Lin, Xinyue Wang, Jing Li, Jie Yin, Fei Gao, Xinyuan Liao, Chenchen Zhang, Qimeng Yin, Chengzhi Zhao, Jinzhong Lin, Yichi Xu, Min Qiu, Dan Luo, Liang Qu

**Author notes:** These authors contributed equally to this work (Yanyan Wang, Liangru Lin, Xinyue Wang, Jing Li, Jie Yin).

## Abstract

Chimeric Antigen Receptor (CAR) T cell therapy has shown promise in treating hematologic malignancies. However, it is limited to individualized cell therapy and faces challenges, including high costs, extended preparation time, and limited efficacy against solid tumors. Here, we generated circular RNAs (circRNAs) encoding Chimeric Antigen Receptor (CAR) transmembrane proteins, referred to as circRNA^CAR^, which mediated remarkable tumor killing in both T cells and macrophages. In addition, macrophages exhibited efficient phagocytosis of tumor cells and pro-inflammatory polarization induced by circRNA^CAR^ *in vitro*. We demonstrated that circRNA^CAR^, delivered with immunocyte-tropic lipid nanoparticles (LNPs), significantly inhibited tumor growth, improved survival rates and induced a pro-inflammatory tumor microenvironment in mice. Importantly, the combination of circRNA^Anti-HER2-CAR^ and circRNA-based cancer vaccines encoding the corresponding transmembrane HER2 antigen, termed circRNA^HER2^, exhibited synergistically enhanced anti-tumor activity. This proof-of-concept study demonstrated that the combination of circRNA-based *in vivo* CAR and vaccines, termed *in vivo* CAR-VAC, holds the potential to become an upgraded off-the-shelf immunotherapy.

## Main Text

Chimeric Antigen Receptor (CAR) T cell therapy is an important method of cancer immunotherapy that has successfully treated hematologic malignancies, particularly B lymphomas^1^. However, CAR-T cell therapy still faces several challenges, including high costs, prolonged preparation time, limited efficacy against solid tumors, and side effects such as the need to clear the marrow, cytokine storm, and on-target, off-tumor killing^2–5^. It is important to address these challenges and improve the effectiveness of CAR-T cell therapy. In comparison to lentiviral^6^ or DNA transposase-mediated delivery^7^, mRNA technology offers a significant safety advantage via not integrating into the genome. Recently, *in vivo* delivery of CAR-encoding mRNAs to T cells or macrophages has been used to produce in vivo CAR-T or CAR-macrophage^7–11^. This innovative approach has shown efficacy in treating hematologic malignancies and myocarditis in mouse models, offering an off-the-shelf immunotherapy strategy with immense potential for therapeutic applications^7–11^. The use of mRNA technology addresses the safety concern of genomic integration and enables broader and more accessible application of CAR-T therapy in diverse clinical scenarios.

Unlike the linear conformation of mRNA, circular RNAs (circRNAs) are covalently closed circular RNA molecules that are more stable than linear mRNAs in mammalian cells^12–14^. Therefore, circRNA-based *in vivo* CAR might enable higher and more durable CAR proteins on the cell membrane of functional immune cells than mRNAs, holding the potential to generate more effective anti-tumor immunity. Recent studies have also shown that the combination therapy of adoptive CAR-T therapy and vaccination could enhance anti-tumor immunity^15–17^. This may occur through further *in vivo* activation of reinfused CAR-T cells via interaction between antigen-presenting cells (APCs) expressing transmembrane tumor antigens and reinfused CAR-T cells expressing CAR molecules against the corresponding antigens^15–17^.

Recently, circRNA vaccines have been developed to elicit effective both B cell and T cell immune responses in mice and rhesus macaques^14^. Currently, therapeutic antibodies were widely used in treating cancers, such as anti-PD1/PD-L1 antibodies and anti-HER2 antibodies, while the anti-tumor activity of antibodies elicited by cancer vaccines, especially the antibody-dependent cellular cytotoxicity (ADCC) or antibody-dependent cellular phagocytosis (ADCP) effects still remain to be clarified^18,19^. Therefore, it is tempting to explore whether circRNA vaccines could synergistically boost the anti-tumor immunity of circRNA-based *in vivo* CAR. Given the potential superiority of circRNAs in generating CAR molecules and producing transmembrane tumor antigens *in vivo*, we attempted to develop the combination immunotherapy between the circRNA-based *in vivo* CAR and cancer vaccines, termed *in vivo* CAR-VAC, aiming to synergistically enhance the anti-tumor activity of *in vivo* CAR via vaccine boosting.

### CircRNA^CAR^ efficiently expressed Chimeric Antigen Receptor (CAR) proteins in both T cells and macrophages

We employed the Group I Intron self-splicing strategy to produce circRNAs that encode anti-HER2 CAR transmembrane proteins, designated as circRNA^CAR^ (Fig. 1a)^14,20^. Subsequently, we utilized high-performance liquid chromatography (HPLC) to obtain high-purity circRNA^CAR^ following RNase R treatment. To assess the translation efficacy of circRNA^CAR^, we transfected the prepared circRNA^CAR^ into HEK293T cells. Western blot revealed clear CAR protein expression (Fig. 1b), and flow cytometry analysis confirmed the transmembrane expression of CAR proteins (Fig. 1c).

**Fig. 1.**
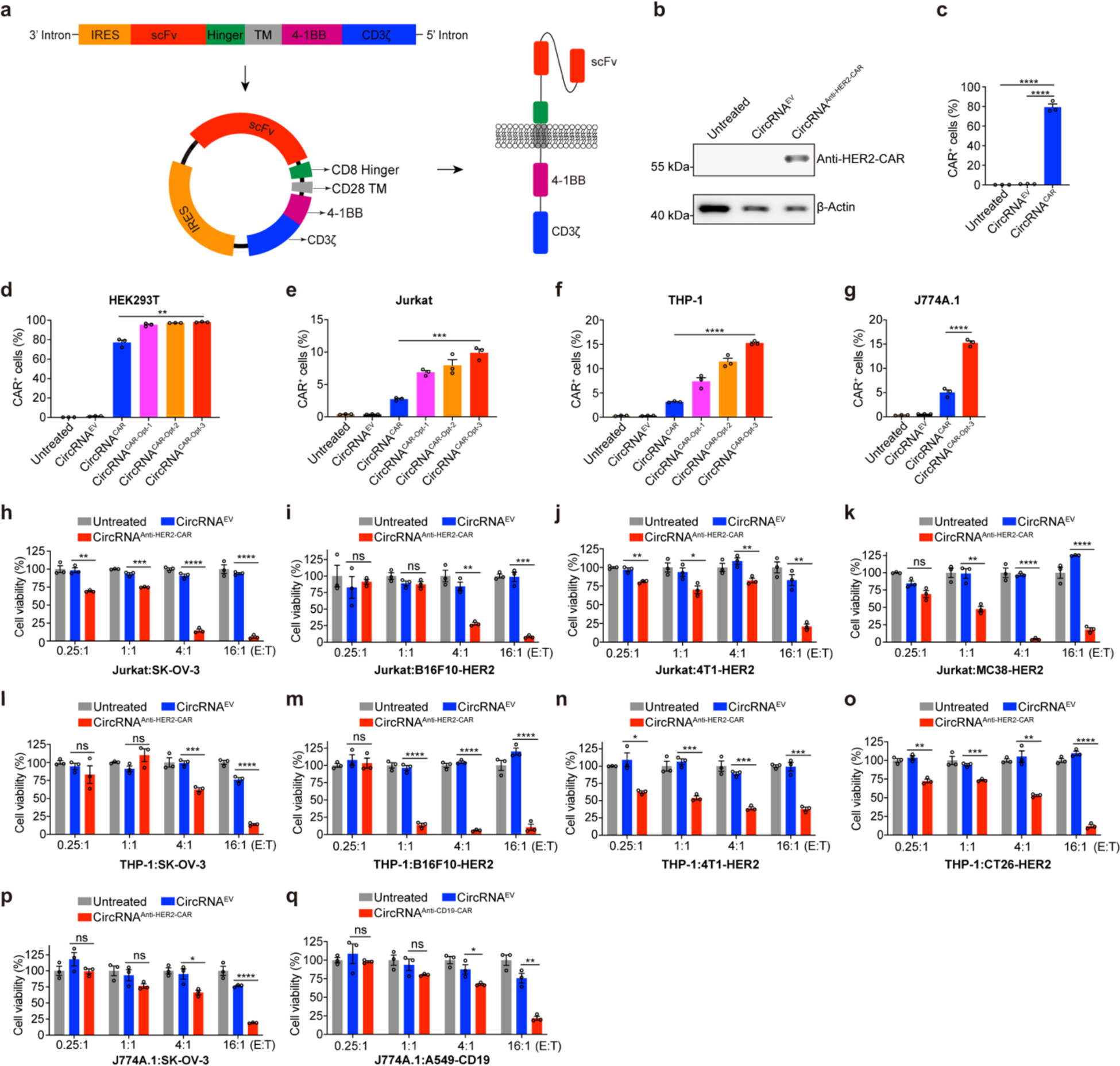
CircRNA^CAR^ efficiently expressed CAR proteins in both T cells and macrophages, and mediated remarkable tumor killing. **a**, Schematic diagram of circRNA^CAR^ circularization by Group I Intron autocatalysis. **b**, **c**, Detecting the expression of CAR proteins in HEK293T cells after circRNA^CAR^ transfection via western blot (**b**) or flow cytometry (**c**). **d**-**g**, Optimization of circRNAs encoding CAR in HEK293T cells (**d**), Jurkat cells (**e**), THP-1 cells (**f**) and J774A.1 cells (**g**). **h**-**k**, Killing effects of Jurkat cells transfected with circRNA^Anti-HER2-CAR^ against SK-OV-3 (**h**), B16F10-HER2 (**i**), 4T1-HER2 (**j**) and MC38-HER2 (**k**) tumor cells. **l**-**o**, Killing effects of THP-1 cells transfected with circRNA^Anti-HER2-CAR^ against SK-OV-3 (**l**), B16F10-HER2 (**m**), 4T1-HER2 (**n**) and CT26-HER2 (**o**) tumor cells. **p**, Killing effects of J774A.1 cells transfected with circRNA^Anti-HER2-CAR^ against SK-OV-3 tumor cells. **q**, Killing effects of J774A.1 cells transfected with circRNA^Anti-CD19-CAR^ against A549-CD19 tumor cells. Data were presented as mean ± S.E.M (n = 3). An unpaired two-sided Student’s t test was performed for comparison, as indicated; *p < 0.05; **p < 0.01; ***p < 0.001; ****p < 0.0001; ns, not significant.

However, while circRNA^CAR^ expressed high levels of CAR proteins in HEK293T cells (Fig. 1d), it produced CAR proteins at much lower levels in both T cells and macrophages (Extended Data Fig. 1a, b). Therefore, we further optimized the protein-encoding sequence of circRNA^CAR^ based on the degeneracy of codons in mammalian cells, aiming to increase its expression in immune cells. We screened out several circRNAs that enable higher expression of CAR proteins in both T cells and macrophages (Fig. 1d-g). Among these optimized circRNAs, circRNA^CAR-opt-3^ exhibits the most efficient expression of CAR proteins (Fig. 1d-g). Therefore, we used the circRNA^CAR-opt-3^ for the following research.

### CircRNA^CAR^ mediated effective tumor killing in both T cells and macrophages

Next, we investigated whether T cells or macrophages transfected with circRNA^CAR^ exhibited cytotoxic effects on the targeted tumor cells. The Jurkat cells, a kind of immortalized human T cells, THP-1 cells, a kind of immortalized human macrophages or J774A.1, a kind of immortalized murine macrophages cells, were transfected with circRNA^Anti-HER2-CAR^ and then co-cultured with various targeted tumor cells, respectively. Subsequently, the tumor-killing effects were detected through luciferase assay, as the above tumor cells stably expressed firefly luciferase. The results of luciferase assay showed that Jurkat cells transfected with circRNA^Anti-HER2-CAR^ could significantly kill multiple targeted tumor cells, including human SK-OV-3 cells (characterized by high expression of HER2), and cancer cell lines ectopically expressing human HER2 including B16F10-HER2 cells, 4T1-HER2 cells, MC38-HER2 cells and CT26-HER2 cells, in an effector:target (E:T) ratio-dependent manner (Fig. 1h-k, and Extended Data Fig.1c). Similarly, circRNA^Anti-HER2-CAR^ also mediated efficient tumor killing in THP-1 or J774A.1 macrophages in an E:T ratio-dependent manner (Fig. 1l-p, and Extended Data Fig. 1d, e). In addition to the circRNA encoding anti-HER2 CAR proteins, we also tested the circRNAs encoding anti-CD19 CAR proteins. We found that THP-1 cells or J774A.1 cells transfected with circRNA^Anti-CD19-CAR^ could also significantly kill human A549 lung carcinoma cells ectopically expressing human CD19 antigens, in an E:T ratio-dependent manner (Fig. 1q and Extended Data Fig. 1f). These results demonstrated that the circRNA^CAR^ could mediated efficient *in vitro* tumor-killing activity in both T cells and macrophages.

### Macrophages exhibited efficient tumor phagocytosis and pro-inflammatory polarization induced by circRNA^CAR^

Macrophages plays a pivotal role in phagocytosis, a process crucial for combating various threats, including tumors^21^. Previous studies have illuminated the role of adeno-virally transduced CAR-Macrophages in mediating tumor phagocytosis^21^. To explore the potential of circRNA^CAR^ in orchestrating tumor phagocytosis by macrophages, the THP-1 or J774A.1 cells was transfected with circRNA^CAR^, and then co-cultured with various targeted tumor cells, respectively. Subsequently, the tumor phagocytosis was detected through flow cytometry, as these tumor cells stably expressed enhanced green fluorescent protein (EGFP) reporter. The flow cytometry analysis unveiled that circRNA^CAR^ mediated significant phagocytosis of diverse cancer cells in both THP-1 and J774A.1 macrophasges in an E:T ratio-dependent manner (Fig. 2a-g, and Extended Data Fig. 2a).

**Fig. 2.**
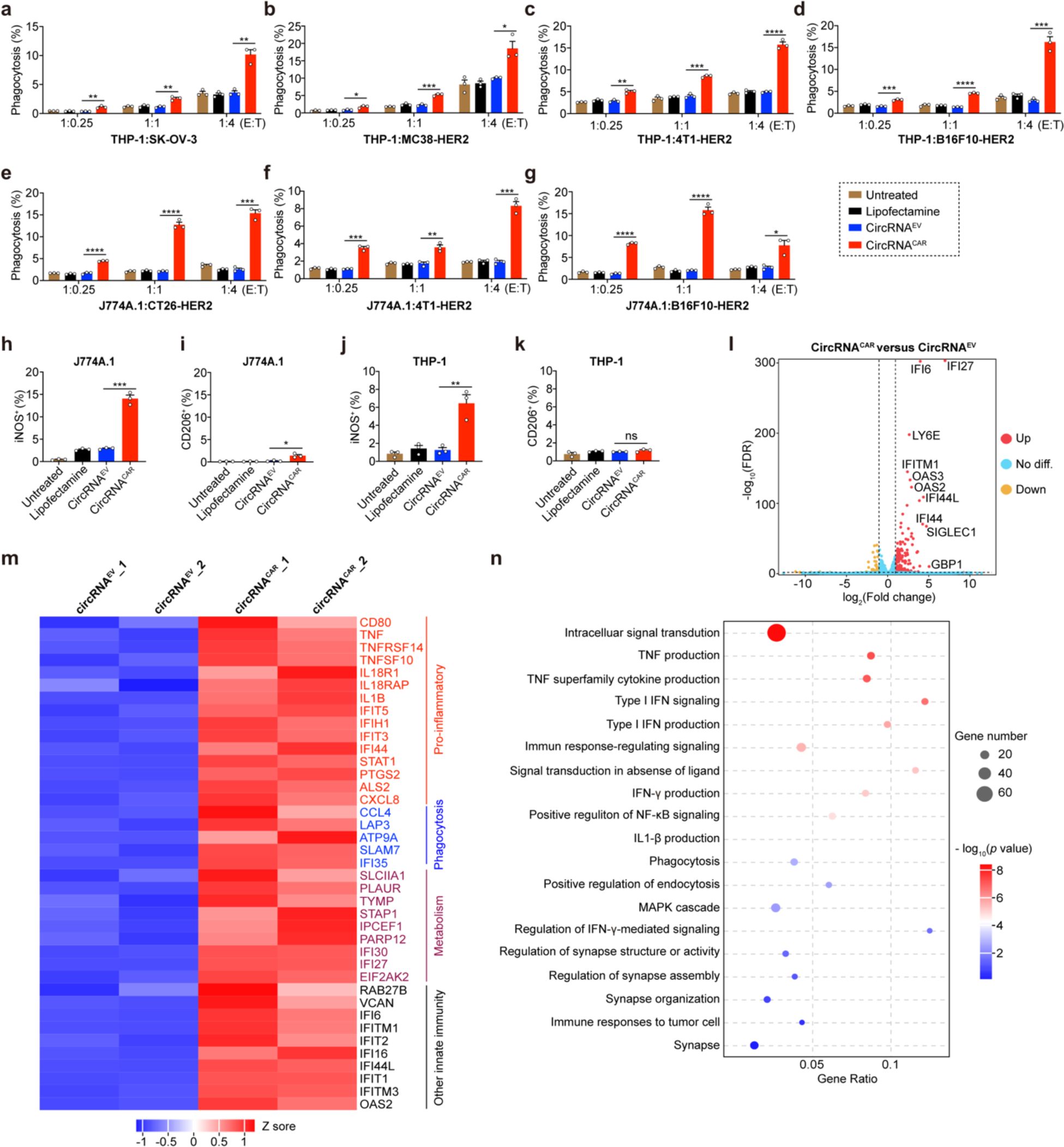
Macrophages exhibited efficient tumor phagocytosis and pro-inflammatory polarization induced by circRNA^CAR^. **a**-**d**, Phagocytosis of THP-1 cells transfected with circRNA^CAR^ against SK-OV-3 (**a**), MC38-HER2 (**b**), 4T1-HER2 (**c**) and B16F10-HER2 (**d**) tumor cells. **e**-**g**, Phagocytosis of J774A.1 cells transfected with circRNA^CAR^ against CT26-HER2 (**e**), 4T1-HER2 (**f**) and B16F10-HER2 (**g**) tumor cells. **h**, **i**, The effects of circRNA^CAR^ on the expression of iNOS (**h**) or CD206 (**i**) in J774A.1 cells. The iNOS or CD206 served as the markers of M1 or M2 polarization state, respectively. **j**, **k**, The effects of circRNA^CAR^ on the expression of iNOS (**j**) or CD206 (**k**) in THP-1 cells. **l**, Volcano plot of differentially expressed genes in the comparison between circRNA^EV^ and circRNA^CAR^ group in THP-1 cells**. m**, Heatmap of gene expression patterns in the comparison between circRNA^EV^ and circRNA^CAR^ group in THP-1 cells (n = 2). **n**, Bubble chart of relevant biological processes involved in the CAR group compared to the EV group through GO analysis (n = 2). The size of the bubbles represented the number of genes. In **a**-**k**, data were presented as mean ± S.E.M (n = 3). An unpaired two-sided Student’s t test was conducted for comparison, as indicated; *p < 0.05; **p < 0.01; ***p < 0.001; ****p < 0.0001; ns, not significant.

In tumor microenvironment, macrophages could be polarized to two opposite states, pro-inflammatory state or anti-inflammatory state ^22^. Subsequently, we tested whether the circRNA^CAR^ could affect the polarization of macrophages. The J774A.1 or THP-1 macrophages were transfected with circRNA^CAR^, respectively. 48 hours later, the macrophages were collected for flow cytometry analysis. The results showed that the circRNA^CAR^-transfected J774A.1 cells demonstrated elevated levels of inducible nitric oxide synthase (iNOS), indicative of the pro-inflammatory phenotype (Fig. 2h). Notably, there was no significant increase in CD206, a marker associated with the anti-inflammatory phenotype (Fig. 2i). Similarly, the circRNA^CAR^-transfected THP-1 cells also exhibited the pro-inflammatory phenotype, but not the anti-inflammatory phenotype (Fig. 2j, k).

To further verify the impact of circRNA^CAR^ on the polarization state of macrophages, we collected the THP-1 cells transfected with PBS, circRNA^EV^ or circRNA^CAR^ for transcriptome-wide RNA-sequencing (RNA-seq) analysis. The results of differential genes expression analysis and cluster analysis showed that circRNA^CAR^ group exhibited upregulation of a series of genes related pro-inflammatory function, phagocytic capability, metabolism and other immune responses of macrophages in comparison with circRNA^EV^ or PBS group (Fig. 2l, m and Extended Data Fig. 2b, c). The pathway enrichment analysis showed that the circRNA^CAR^ group exhibited enhanced intracellular signal transduction and several enriched inflammatory-associated pathways such as TNF production, IFN-γ production, NF-κB signaling and MAPK cascade signaling (Fig. 2n and Extended Data Fig. 2d). Collectively, these results suggested that circRNA^CAR^ held the potential to induce a pro-inflammatory M1 polarization in macrophages, which provided novel insights into the immunomodulatory capabilities of circRNA^CAR^.

### Screening for immune-tropic lipid nanoparticles that efficiently delivered circRNAs into the immune cells in mice

It is crucial to efficiently deliver circRNA^CAR^ into T cells and macrophages *in vivo*. The spleen, predominantly composed of immune cells such as T cells, B cells, and macrophages, is an important immune organ for the targeted delivery of circRNA^CAR^ using LNPs^23–27^. Therefore, we synthesized various cationic lipids for LNPs by altering the head and tail groups. We screened out an LNP composition that contains four key components: the specific cationic lipids, cholesterol, DSPC and PEG2000-DMG, which efficiently targeted the spleen (Fig. 3a). Then, the circRNA^Luciferase^, encoding the firefly luciferase, was encapsulated within this LNP. The average size of the LNP-circRNA^Luciferase^ formulation is approximately 110 nm (Fig. 3b). The LNP-circRNA^Luciferase^ formulation also exhibited positive electrical properties, as demonstrated by a zeta potential of approximately 8 mV (Fig. 3c). To investigate the *in vivo* targeting effects, the LNP-circRNA^Luciferase^ was intravenously injected into BALB/c mice. Six hours later, we assessed the tissue distribution of LNP-circRNA^Luciferase^ using three-dimensional *in vivo* and *ex vivo* luciferase imaging (Fig. 3d, e). The imaging results showed that the luciferase was mainly concentrated in the spleen, but not the liver or lung, indicating the immunocyte-tropic feature of this LNP. In addition, the tumor microenvironment contained substantial tumor-associated macrophages, T cells, and dendritic cells^28^. And then the LNP-circRNA^Luciferase^ was subsequently locally injected into the tumors of subcutaneous tumor-bearing mice. *In vivo* luciferase imaging results showed that the luciferase was mainly concentrated within the local tumor region (Fig. 3f). Collectively, these results demonstrated the capability of LNP-circRNA for immune-tropic delivery *in vivo*.

**Fig. 3.**
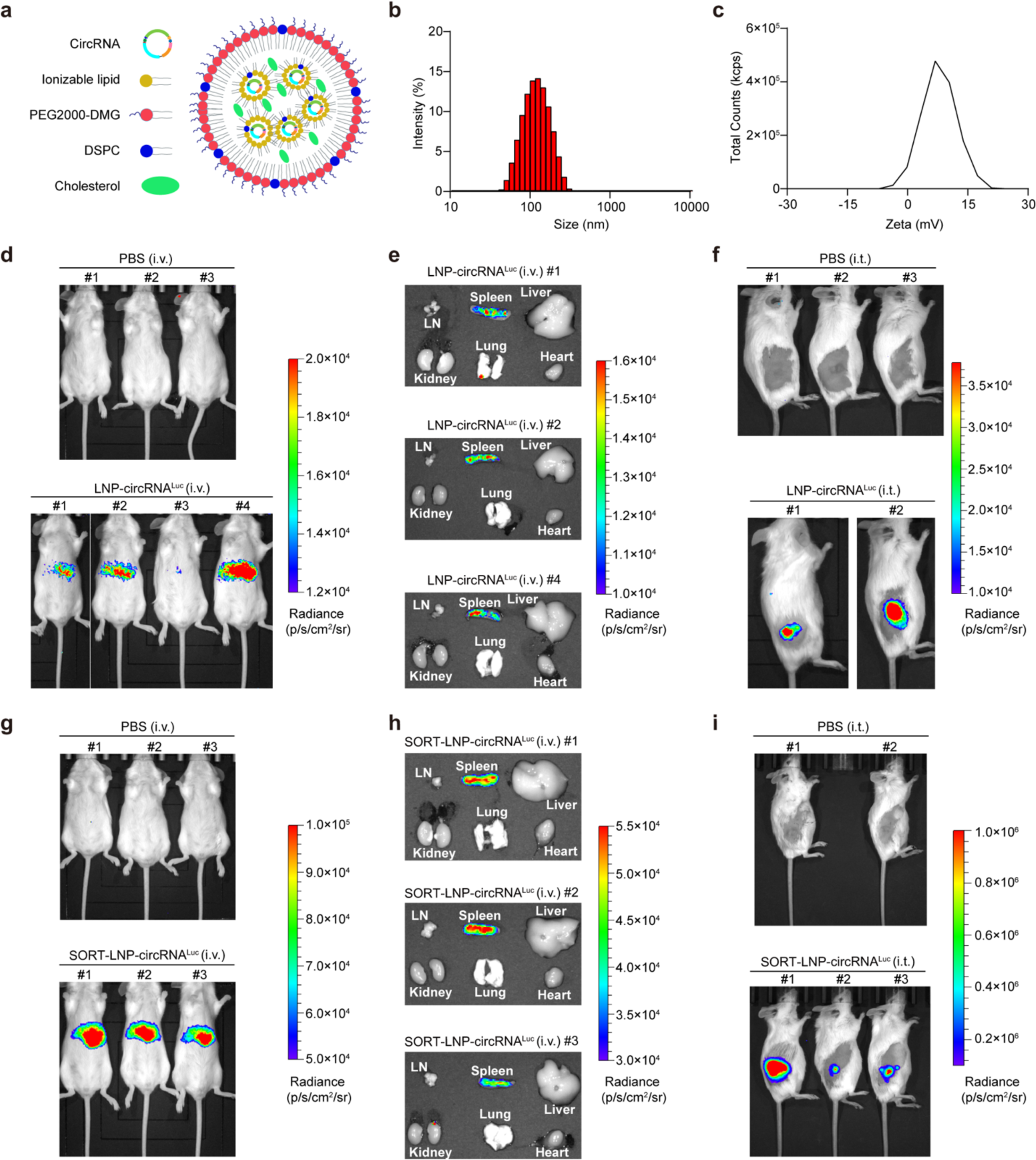
Screening for immune-tropic lipid nanoparticles that efficiently delivered circRNAs into the immune cells in mice. **a**, Schematic representation of the LNP-circRNA complex. **b**, The size distribution of LNP-circRNA^Luciferase^ complex. **c**, The zeta potential of LNP-circRNA^Luciferase^ complex. **d**, **e**, Bioluminescence imaging *in vivo* (**d**) or *ex vivo* (**e**) of BALB/c mice intravenously injected with PBS or LNP-circRNA^Luciferase^ (n = 3). **f**, Bioluminescence imaging of BALB/c mice intratumorally injected with PBS or LNP-circRNA^Luciferase^ (n = 2 or 3). **g**, **h**, Bioluminescence imaging *in vivo* (**g**) or *ex vivo* (**h**) of BALB/c mice intravenously injected with PBS or SORT-LNP-circRNA^Luciferase^ (n = 3). **i**, Bioluminescence imaging of BALB/c mice intratumorally injected with PBS or SORT-LNP-circRNA^Luciferase^ (n = 2 or 3).

Besides, we also tested the Selective ORgan Targeting (SORT) approach, a previously reported method where lipid nanoparticles are engineered with complementary SORT molecules for precise targeting of extrahepatic tissues^23^. This technique employs negatively charged SORT-LNP for targeted spleen delivery. To evaluate the effect of *in vivo* targeting, we intravenously administered SORT-LNP-circRNA^Luciferase^ into BALB/C mice. Six hours later, we analyzed the tissue distribution using both *in vivo* and *ex vivo* luciferase imaging. The imaging results revealed a predominant luciferase expression in the spleen, indicating the immune-tropic specificity of negatively charged SORT-LNP (Fig. 3g, h). Finally, we also tested the intratumoral injection of SORT-LNP-circRNA^Luciferase^ in tumor-bearing mice, followed by the *in vivo* luciferase imaging. The imaging results elucidated a localized high concentration in tumor region (Fig. 3i), which was consistent with the above LNP we initially prepared.

### CircRNA^CAR^ efficiently inhibited tumor growth and improved survival time in mice

To test whether the circRNA^CAR^ can inhibit tumor growth *in vivo*, we firstly used the immune-competent BALB/c mice with CT26-HER2 colorectal carcinomas (Fig. 4a). When the subcutaneous CT26-HER2 tumor volume reached approximately 50∼80 mm^3^, the tumor-bearing mice were randomly grouped and intravenously administrated three times with PBS, 15 μg of SORT-LNP-circRNA^EV^ or 15 μg of SORT-LNP-circRNA^CAR^, and then the tumor volume of grouped mice was monitored (Fig. 4a). The result of tumor volume detection showed that SORT-LNP-circRNA^CAR^ significantly inhibited CT26-HER2 tumor growth, in comparison with PBS or SORT-LNP-circRNA^EV^ groups (Fig. 4b).

**Fig. 4.**
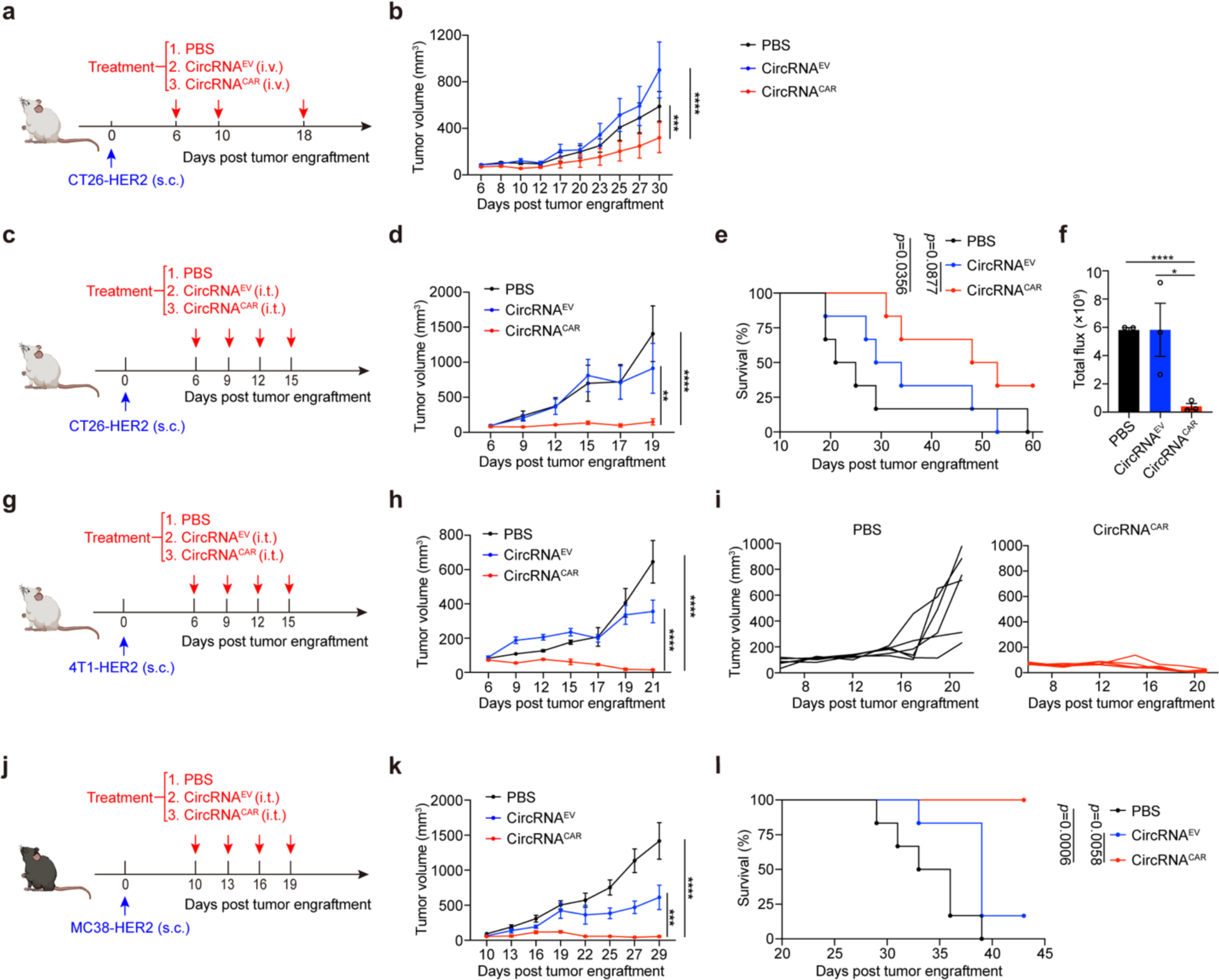
CircRNA^CAR^ efficiently inhibited tumor growth and improved survival time in mice. **a**, Schematic illustration of PBS, SORT-LNP-circRNA^EV^ or SORT-LNP-circRNA^CAR^ treatment in CT26-HER2 tumor model via intravenous administration. **b**, Tumor growth curves of CT26-HER2 tumor-bearing mice treated with intravenous administration of PBS, SORT-LNP-circRNA^EV^ or SORT-LNP-circRNA^CAR^. **c**, Schematic illustration of PBS, LNP-circRNA^EV^ or LNP-circRNA^CAR^ treatment in CT26-HER2 tumor model via intratumoral administration. **d**, **e**, Tumor growth curves (**d**) and survival curves (**e**) of CT26-HER2 tumor-bearing mice treated with PBS, LNP-circRNA^EV^ or LNP-circRNA^CAR^ via intratumoral administration. **f**, *In vivo* bioluminescence imaging of CT26-HER2 tumor-bearing mice after the treatment of PBS, LNP-circRNA^EV^ or LNP-circRNA^CAR^ via intratumoral administration. **g**, Schematic illustration of PBS, LNP-circRNA^EV^ or LNP-circRNA^CAR^ treatment in 4T1-HER2 tumor model via intratumoral administration. **h**, Tumor growth curves of 4T1-HER2 tumor-bearing mice treated with PBS, LNP-circRNA^EV^ or circRNA^CAR^ via intratumoral administration. **i**, Tumor growth curves of individual 4T1-HER2 tumor-bearing mice treated with PBS, or LNP-circRNA^CAR^ via intratumoral administration. **j**, Schematic illustration of PBS, LNP-circRNA^EV^ or LNP-circRNA^CAR^ treatment in MC38-HER2 tumor model via intratumoral administration. **k**, **l**, Tumor growth curves (**k**) and survival curves (**l**) of MC38-HER2 tumor-bearing mice treated with PBS, LNP-circRNA^EV^ or LNP-circRNA^CAR^ via intratumoral administration. Data were represented as the mean ± S.E.M. In **b**, **d**, **h** and **k**, the tumor growth curves were calculated by two-way ANOVA analysis (n = 6). In **e** and **l**, the survival curves were calculated by Kaplan-Meier simple survival analysis (n = 6). In **f**, an unpaired two-sided Student’s t test was conducted for comparison, as indicated. *p < 0.05; **p < 0.01; ***p < 0.001; ****p < 0.0001; ns, not significant.

Apart from the intravenous administration, we also tested the intratumoral administration in immune-competent BALB/c mice with subcutaneous CT26-HER2 colonic colorectal carcinomas (Fig. 4c). Similarly, when the subcutaneous CT26-HER2 tumor volume reached approximately 50∼80 mm^3^, the tumor-bearing mice were randomly grouped and intratumorally injected four times at a three-day interval with PBS, 15 μg of LNP-circRNA^EV^ or 15 μg of LNP-circRNA^CAR^ (Fig. 4c). We found that LNP-circRNA^CAR^ could significantly also significantly reduced tumor burden and prolonged survival time, in comparison with PBS or LNP-circRNA^EV^ groups (Fig. 4d, e). And the *in vivo* luciferase imaging results also showed that LNP-circRNA^CAR^ tumor burden of LNP-circRNA^CAR^-treated groups was significantly lower than that of PBS or LNP-circRNA^EV^-treated groups (Fig. 4f). Similarly, in the 4T1 breast carcinomas-bearing BALB/c mice (Fig. 4g), the intratumoral administration of LNP-circRNA^CAR^ significantly inhibited tumor growth and even achieved the complete regression of subcutaneous carcinomas, in comparison with the PBS or LNP-circRNA^EV^ groups (Fig. 4h, i). As for the subcutaneous MC38-HER2 colonic colorectal carcinomas (Fig. 4j), the intratumoral administration of LNP-circRNA^CAR^ also significantly inhibited tumor growth and increased the survival time of tumor-bearing mice, in comparison with PBS or LNP-circRNA^EV^ groups (Fig. 4k, l).

Collectively, these results demonstrated that the circRNA-based *in vivo* CAR exerted promising efficacy in treating solid tumors.

### CircRNA^CAR^ reshaped the tumor microenvironment to a pro-inflammatory state in mice

Solid tumors are often refractory partly due to the immunosuppressive tumor microenvironment^29^. Therefore, we further investigated the effects of circRNA^CAR^ on the tumor microenvironment of tumor-bearing mice. At the end of experiments via intravenous administration of PBS, circRNA^EV^ or circRNA^CAR^, the mouse spleen and tumor tissues were collected for detecting the infiltration of immune cells via flow cytometry analysis. The flow cytometry results showed that the circRNA^CAR^ group exhibited significant increase in the proportion of CD8^+^ T cells, CD8^+^ effector memory T cells (Tem), central memory T cells (Tcm) and CD4^+^ T cells, but a significant decrease in the proportion of Treg cells in the spleen, compared to the PBS or circRNA^EV^ group (Fig. 5a). As for tumor tissues, we also found that, the PBS or circRNA^EV^ group, circRNA^CAR^ group exhibited increased proportion of tumor-infiltrating CD45^+^ cells, CD8^+^ T cells and CD4^+^ T cells, but a decreased proportion of tumor-infiltrating Treg cells (Fig. 5b). The flow cytometry results also showed that the circRNA^CAR^-treated mice exhibited significantly increase the proportion of pro-inflammatory macrophages in the tumor microenvironment in comparison with the PBS or circRNA^EV^ groups (Fig. 5b).

**Fig. 5.**
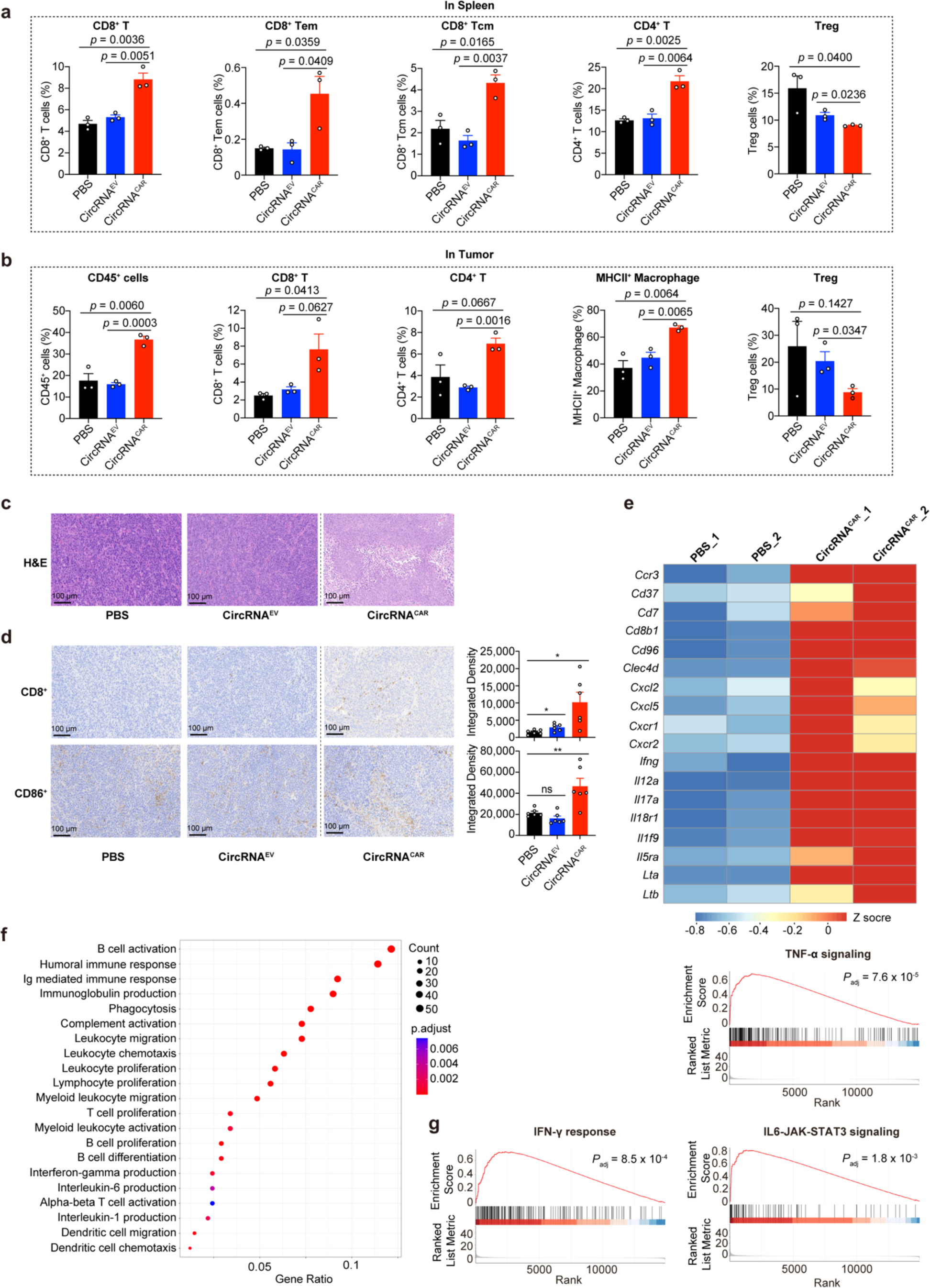
CircRNA^CAR^ reshaped the tumor microenvironment to a pro-inflammatory state in mice. **a**, **b**, Flow cytometric analysis of spleen cells (**a**) and tumor-infiltrating lymphocytes (**b**) extracted from CT26-HER2 tumor-bearing mice treated with intravenous administration of PBS, SORT-LNP-circRNA^EV^ or SORT-LNP-circRNA^CAR^. CD8^+^ T cells, CD8^+^ Tem cells, CD8^+^ Tcm cells, CD4^+^ T cells or MHCII^+^ Macrophages were gated from CD45^+^ cell population, and Treg cells were gated from CD4^+^ T cell population. **c**, **d**, H&E staining of tumor tissue sections (**c**) and IHC staining of infiltrated CD8^+^ cells or CD86^+^ cells in tumor tissue (d) obtained from CT26-HER2 tumor-bearing mice treated with PBS, LNP-circRNA^EV^ or LNP-circRNA^CAR^ via intratumoral administration. The integrated density of IHC staining was quantified using ImageJ software. **e**, Heatmap of gene expression patterns of immune cells extracted from tumor tissues in the comparison between PBS and circRNA^CAR^ group via intratumoral administration (n = 2). **f**, Bubble chart of relevant biological processes involved in the circRNA^CAR^ group compared to the PBS group via intratumoral administration through GO analysis (n = 2). The size of the bubbles represented the number of genes. **g**, Gene Set Enrichment Analysis (GSEA) showing enriched pathways in the immune cells extracted from tumor tissues in the comparison between PBS and circRNA^CAR^ group via intratumoral administration (n = 2). In **a**, **b** and **d**, an unpaired two-sided Student’s t test was conducted for comparison, as indicated; *p < 0.05; **p < 0.01; ***p < 0.001; ****p < 0.0001; ns, not significant. Each symbol represents an individual mouse.

Besides, we also harvested the tumor tissue specimens from the treated mice for Hematoxylin and eosin (H&E) staining and Immunohistochemistry IHC staining. The results of H&E staining showed that the circRNA^CAR^-treated group exhibited significant increase in tumor cell death with enlargement of tumor cell gaps, pale cytoplasm, cell disintegration, and loss of structure, while the PBS or circRNA^EV^ groups exhibited diffuse growth and dense arrangement in both the CT26-HER2 and MC38-HER2 and colorectal carcinoma-bearing mice (Fig. 5c and Extended Data Fig. 3a, b). The results of IHC staining demonstrated an increase in the infiltration of both CD8^+^ T cells and pro-inflammatory CD86^+^ cells in the circRNA^CAR^ groups, compared to the PBS or circRNA^EV^ groups in both MC38-HER2 and CT26-HER2 colorectal carcinoma-bearing mice (Fig. 5d and Extended Data Fig. 3c, d).

To further explore the impact of circRNA^CAR^ on tumor microenvironment, we isolated the immune cells from tumor tissues via density gradient centrifugation and performed RNA-seq analysis. The differential gene expression analysis showed that the circRNA^CAR^ group exhibited upregulation of genes pro-inflammatory cytokines, chemokines and chemokine receptors, in comparison with the circRNA^EV^ or PBS group, suggesting the infiltration of immune cells in the tumor microenvironment (Fig. 5e and Extended Data Fig. 3e). The pathway enrichment analysis showed that the circRNA^CAR^ group exhibited several enriched pathways related to T cell proliferation, Dendritic cell migration, phagocytosis and pro-inflammatory responses such as IFN-γ signaling, TNF-α signaling, IL6-JAK-STAT3 signaling, in comparison with the PBS or LNP-circRNA^EV^ groups (Fig. 5f, g and Extended Data Fig. 3f, g). Notably, we found that multiple pathways related to B cell immunity, such as B cell activation, B cell proliferation, Immunoglobulin mediated immune response, Immunoglobulin production and complement activation, were significantly enriched in the circRNA^CAR^ group, in comparison with the PBS or LNP-circRNA^EV^ group (Fig. 5f and Extended Data Fig. 3f). This finding suggested that the antibodies might play an important role in the anti-tumor immunity of *in vivo* CAR therapy, warranting further investigation. In summary, these results demonstrated that circRNA^CAR^ could markedly reshape tumor microenvironment to a pro-inflammatory state in tumor-bearing mouse model, which was tended to sensitizing immunotherapy.

### CircRNA vaccine synergistically boosted the anti-tumor activity of *in vivo* CAR therapy via antibody-mediated cellular cytotoxicity

Recent reports proposed that vaccines might further enhance the therapeutic efficacy of CAR-T adoptive cell therapy in mouse tumor models^15–17^. Notably, the targeted proteins of the CAR molecules align with the antigens of the cancer vaccine^16^, which were designed to induce antigen spreading through the reactivation of the infused CAR-T cells by the corresponding antigens of cancer vaccine^17^. The above RNA-seq results suggested that the anti-tumor effects of circRNA^CAR^ therapy might be related to B cell responses (Fig. 5f and Extended Data Fig. 3f). Additionally, antibodies could promote the phagocytosis or killing effects of macrophages or NK cells through antibody-dependent cellular cytotoxicity (ADCC) and antibody-dependent cellular phagocytosis (ADCP)^18,19^. Therefore, we aimed to investigate whether circRNA vaccines could synergistically boost the anti-tumor immunity of the circRNA-based *in vivo* CAR, in the manner of ADCC or ADCP effects.

As the circRNA^Anti-HER2-CAR^ encoded anti-HER2 CAR molecules that targeting human HER2 proteins, we developed a circRNA vaccine that encoded a human HER2 fusion protein, in which the endocytosis prevention motif (EPM) and ESCRT- and ALIX-binding region (EABR) domain was fused at the C teminal instead of the original intracellular domain of human HER2 protein, aiming to help elicit higher level of HER2-specific antibodies^30^ (Fig. 6a). Therefore, we tested the combination immunotherapy between the circRNA-based *in vivo* CAR and circRNA-based cancer vaccines (*in vivo* CAR-VAC). The circRNA vaccine termed circRNA^VAC^, encoding HER2-EPM-EABR, which could enhance the vesicle secretion of the HER2 antigens by the EPM-EABR motif and improve the humoral immune responses^30^. In HEK293T cells, circRNA^VAC^ exhibited efficient trans-membrane expression of HER2-EPM-EABR fusion protein (Fig. 6b), concomitant with the efficient secretion of vesicles in the supernatant (Fig. 6c).

**Fig. 6.**
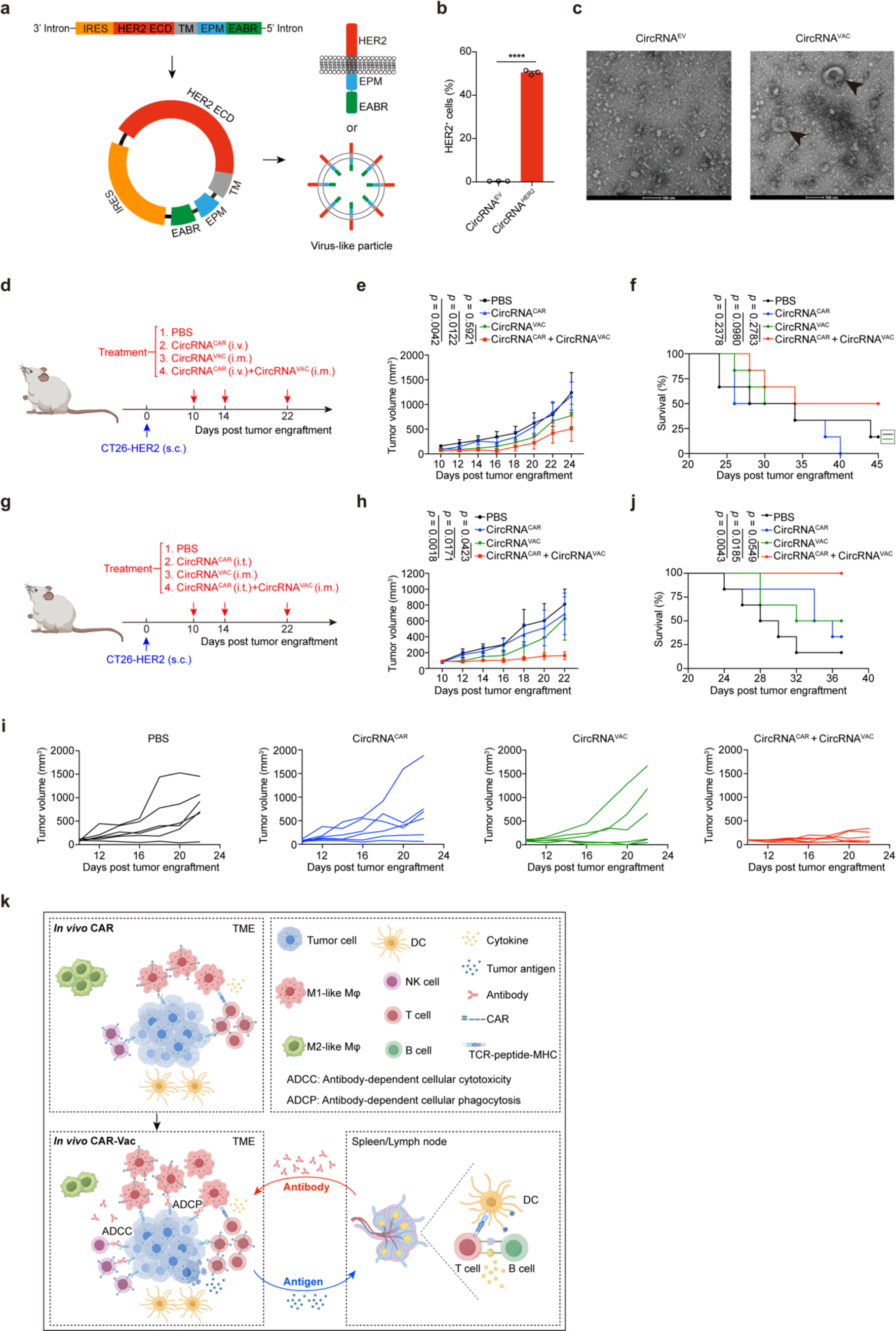
CircRNA vaccine synergistically boosted the anti-tumor activity of in vivo CAR therapy. **a**, Schematic illustration of the circRNA^VAC^ design. EPM, the endocytosis prevention motif; EABR, the ESCRT- and ALIX-binding region domain. **b**, Flow cytometric analysis detecting the translation of circRNA^VAC^ encoding HER2-EPM-EABR fusion proteins in HEK293T cells (n = 3). **c**, HEK293T cells transfected with circRNA^VAC^ secreted vesicles carrying transmembrane HER2 antigens in the supernatant. **d**, Schematic illustration of *in vivo* circRNA^CAR^ (intravenously) therapy combined with circRNA^VAC^ (intramuscularly). **e**, Tumor growth curves of overall CT26-HER2 tumor-bearing mice after receiving PBS (intravenously), circRNA^CAR^ (intravenously), circRNA^VAC^ (intramuscularly), or circRNA^CAR^ (intravenously) plus circRNA^VAC^ (intramuscularly) combined therapy (n = 6). **f**, Tumor Survival curves of CT26-HER2 tumor-bearing mice after receiving PBS (intravenously), circRNA^CAR^ (intravenously), circRNA^VAC^ (intramuscularly), or circRNA^CAR^ (intravenously) plus circRNA^VAC^ (intramuscularly) combined therapy (n = 6). **g**, Schematic illustration of *in vivo* circRNA^CAR^ (intratumorally) therapy combined with circRNA^VAC^ (intramuscularly). **h**, Tumor growth curves of overall CT26-HER2 tumor-bearing mice after receiving PBS (intratumorally), circRNA^CAR^ (intratumorally), circRNA^VAC^ (intramuscularly), or circRNA^CAR^ (intratumorally) plus circRNA^VAC^ (intramuscularly) combined therapy (n = 6). **i**, Tumor growth curves of individual CT26-HER2 tumor-bearing mice treated as indicated in (**h**) (*n* = 6). **j**, Survival curves of CT26-HER2 tumor-bearing mice treated as indicated in (**h**) (*n* = 6). **k**, The potential mechanism diagram of synergistic *in vivo* CAR-VAC immunotherapy. In **b**, data were represented as the mean ± S.E.M; an unpaired two-sided Student’s *t*-test was performed for comparison as indicated in the figure. In **e** and **h**, the data were represented as the mean ± S.E.M, the tumor growth curves were calculated by two-way ANOVA analysis. In **f** and **j**, the survival curves were calculated by Kaplan-Meier simple survival analysis. *p < 0.05; **p < 0.01; ***p < 0.001; ****p < 0.0001; ns, no significant.

To test the effects of *in vivo* CAR-VAC therapy, the C57BL/6 mice with subcutaneous CT26-HER2 colorectal carcinomas, were injected with PBS control (intravenously), 15 μg of circRNA^CAR^ (intravenously), 15 μg of circRNA^VAC^ (intramuscularly), or 15 μg of circRNA^CAR^ (intravenously) plus 15 μg of circRNA^VAC^ (intramuscularly) at 10, 14, 22 days post tumor engraftment (Fig. 6d). The results showed that the combination of the intravenously delivered circRNA^CAR^ and circRNA^HER2^ vaccine showed improved tumor inhibition and enhanced survival time, in comparison with the PBS control, individual circRNA^CAR^ or individual circRNA^VAC^ groups (Fig. 6e, f).

Additionally, we also investigated the efficacy of intratumoral administration, as an alternative to the intravenous administration, for the delivery of circRNA^CAR^ (Fig. 6g). Impressively, the combination of the intratumoral delivered circRNA^CAR^ and circRNA^VAC^ vaccine markedly impeded tumor growth, resulting in a noteworthy enhancement in the survival rate of tumor-bearing mice (Fig. 6h-j). Traditional developing strategies of cancer vaccines have been mainly focusing on the T cell-mediated anti-tumor immunity, but not the B cell-mediated anti-tumor immunity, whereas in this study we found that B cell-mediated immune responses played an important role in tumor suppression. On one hand, circRNA vaccine-elicited tumor-specific antibodies might boost the *in vivo* circRNA^CAR^-mediated tumor killing via ADCC and ADCP effects, beyond the basic CAR-mediated phagocytosis or killing effects in TME (Fig. 6k). On the other hand, in return, the elevated exposure of tumor antigens might further boost the B cell immunity to produce potent tumor-specific antibodies in the germinal centers of tumor-draining lymph node (TDLN) or spleen (Fig. 6k). This potential positive feedback loop between antibody and antigen might partly explain the synergically enhanced anti-tumor immunity of *in vivo* CAR-VAC immunotherapy.

Collectively, these results demonstrated that the circRNA vaccines could synergistically boost the anti-tumor immunity of circRNA-based *in vivo* CAR delivered either intratumorally or intravenously, which holds the potential for an upgraded off-the-shelf *in vivo* CAR-VAC immunotherapy for treating solid tumors.

## Discussion

CircRNA, a covalently closed circular RNA molecule, is highly stable and has been engineered for various biomedical applications, such as RNA editing, RNA vaccines against infectious diseases, cancer vaccines and gene therapy^31–36^. Recent research reported that the circRNA-based TCR-T adoptive cell therapy for treating cytomegalovirus infection ^37^. In this study, we generated circRNA^CAR^ which encoded anti-HER2 CAR transmembrane proteins for *in vivo* CAR immunotherapy to treat solid tumors in mice. Via optimizing the sequence of circRNA^CAR^, we also improved the translation efficiency of circRNA^CAR^ in both T cells and macrophages (Fig. 1d-g).

Subsequently, we demonstrated that circRNA^CAR^-transfected T cells exhibited efficient *in vitro* tumor killing (Fig. 1h-k). Similarly, the macrophages transfected with circRNA^CAR^ also exhibited efficient *in vitro* killing and phagocytosis (Fig. 1 and 2). Notably, we also found that circRNA^CAR^-transfected J774A.1 or THP-1 cells exhibited pro-inflammatory polarization, which provided important insights into the multifaceted immunomodulatory capabilities of the circRNA-based *in vivo* CAR immunotherapy (Fig. 2h-n and Extended Data Fig. 2b-d).

For *in vivo* study, using the immune-tropic lipid nanoparticles (Fig. 3), circRNA^CAR^ was delivered into tumor-bearing mice and effectively suppressed tumor growth and markedly improved the survival rate in multiple mouse models (Fig. 4). Moreover, we also found circRNA^CAR^ could reshape the tumor microenvironment to a highly pro-inflammatory state in both MC38-HER2 and CT26-HER2 colorectal carcinoma-bearing mouse model, facilitating the infiltration of CD8^+^ T cells and pro-inflammatory CD86^+^ cells (Fig. 5, Extended Data Fig. 3). Therefore, this circRNA-based *in vivo* CAR therapy held the potential to sensitize the anti-tumor immunity via combination with other cancer therapies, such as the targeted therapy or immune checkpoint blockade therapies (eg. PD1/PD-L1 antibodies). In comparison with the current adoptive CAR-T cell therapy, this off-the-shelf *in vivo* circRNA^CAR^ therapy potentially had multiple advantages such as (1) lower cost; (2) off-the-shelf therapy; (3) avoiding the risk of genomic integration; (4) without marrow clearance; (5) Redosable.

Previous research reported that mRNA vaccines encoding tumor-associated antigens could help enhance the anti-tumor effects of adoptive CAR-T cell therapy^16^. Up to now, to my knowledge, there are no reports about the combined immunotherapy between *in vivo* CAR therapy and RNA vaccines. More importantly, antibodies play an important role in macrophages or NK cells-mediated phagocytosis or killing via ADCC or ADCP effects^18,19,38,39^. Current studies on the combined therapy of CAR-T adoptive cell therapy and vaccines have only reported on the interactions between CAR-T cells and antigen-presenting cells^17^, while the functions of vaccination-elicited antibodies still remain to be clarified. In this study, we formulated a circRNA-based cancer vaccine encoding human HER2 antigens with the fused intracellular EPM-EABR motif to enhance the level of HER2-specific antibodies^30^. We further demonstrated that this circRNA vaccine could synergistically boost the anti-tumor immunity of circRNA-based *in vivo* CAR delivered either intratumorally or intravenously (Fig. 6). Mechanistically, it was possible that the immune cells expressing CAR functional molecules, such as T cells, macrophages and NK cells, could specifically kill tumor cells, release inflammatory cytokines, reshape the immunosuppressive tumor microenvironment, and promote the infiltration of pro-inflammatory immune cell.

In summary, this proof-of-concept study established a novel combined immunotherapy between *in vivo* CAR and cancer vaccines, which achieved synergistically enhanced anti-tumor effects in multiple mouse models, providing an upgraded off-the-shelf *in vivo* CAR immunotherapy for treating solid tumors in the future.

---

**Extended Data Fig. 1.**
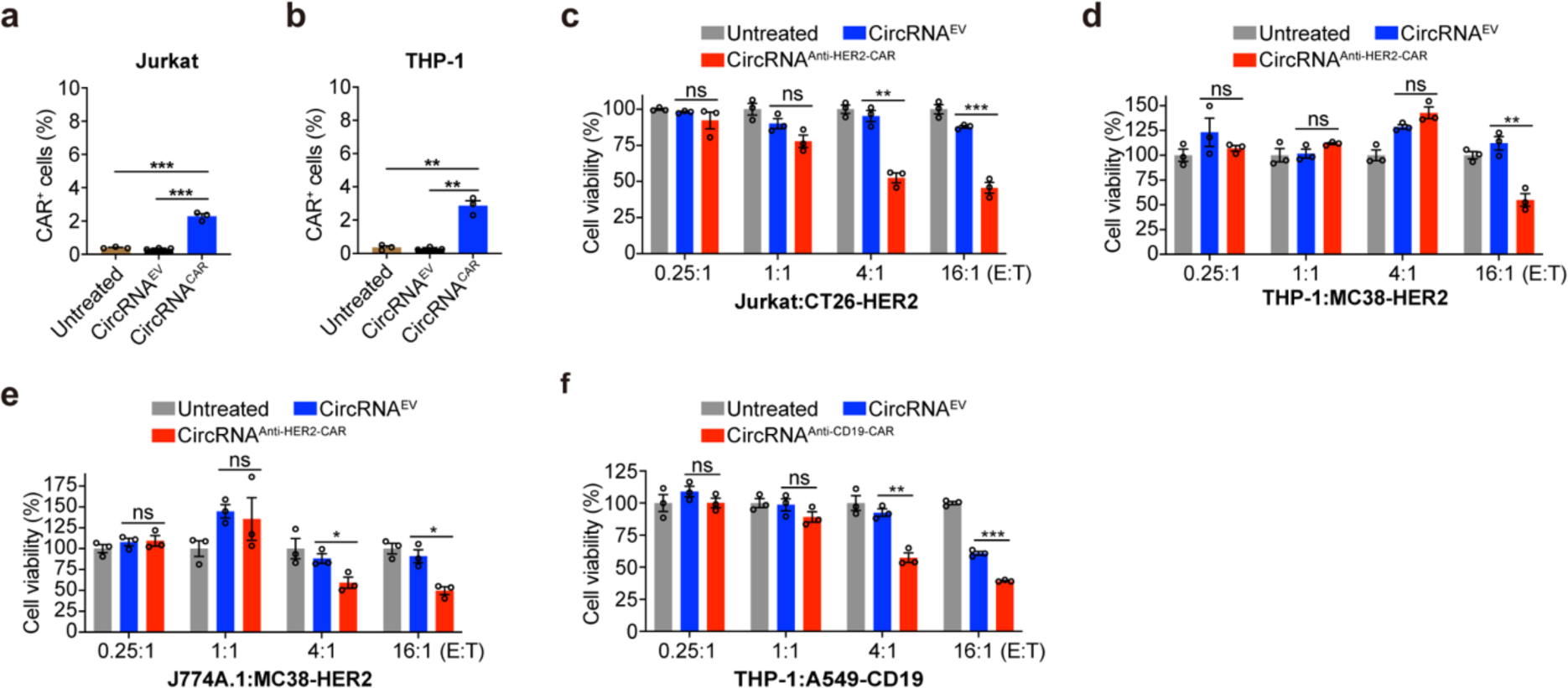
CircRNA^CAR^ expressed CAR proteins in both T cells and macrophages, and mediated remarkable tumor killing. **a**, **b**, Detecting the expression of CAR proteins in both Jurkat and THP-1 cells via flow cytometry. **c**, Killing effects of Jurkat cells transfected with circRNA^Anti-HER2-CAR^ against CT26-HER2 tumor cells. **d**, **e**, Killing effects of THP-1 cells (**d**) and J774A.1 cells (**e**) transfected with circRNA^Anti-HER2-CAR^ against MC38-HER2 tumor cells. **f**, Killing effects of THP-1 cells transfected with circRNA^Anti-CD19-CAR^ against A549-CD19 tumor cells. Data were presented as mean ± S.E.M (n = 3). An unpaired two-sided Student’s t test was performed for comparison, as indicated; *p < 0.05; **p < 0.01; ***p < 0.001; ****p < 0.0001; ns, not significant.

**Extended Data Fig. 2.**
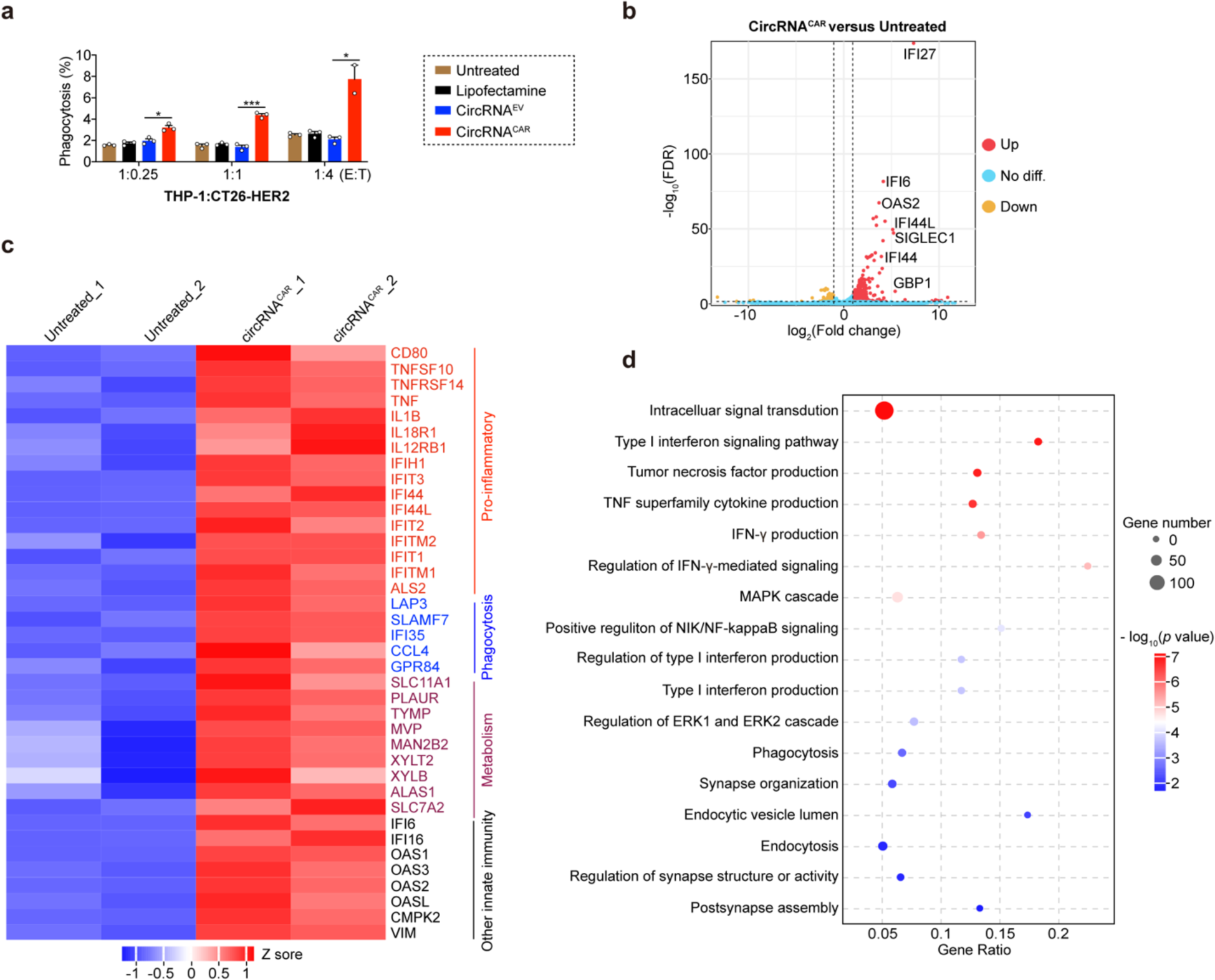
Macrophages exhibited efficient tumor phagocytosis and pro-inflammatory polarization induced by circRNA^CAR^. **a**, Phagocytosis of THP-1 cells transfected with circRNA^CAR^ against CT26-HER2 tumor cells. **b**, Volcano plot of differentially expressed genes in the comparison between PBS and circRNA^CAR^ group in THP-1 cells. **c**, Heatmap of gene expression patterns in the comparison between PBS and circRNA^CAR^ group in THP-1 cells (n = 2). **d**, Bubble chart of relevant biological processes involved in the circRNA^CAR^ group compared to the PBS group through G analysis (n = 2). The size of the bubbles represented the number of genes. In **a**, data were presented as mean ± S.E.M (n = 3). An unpaired two-sided Student’s t test was conducted for comparison, as indicated; *p < 0.05; **p < 0.01; ***p < 0.001; ****p < 0.0001; ns, not significant.

**Extended Data Fig. 3.**
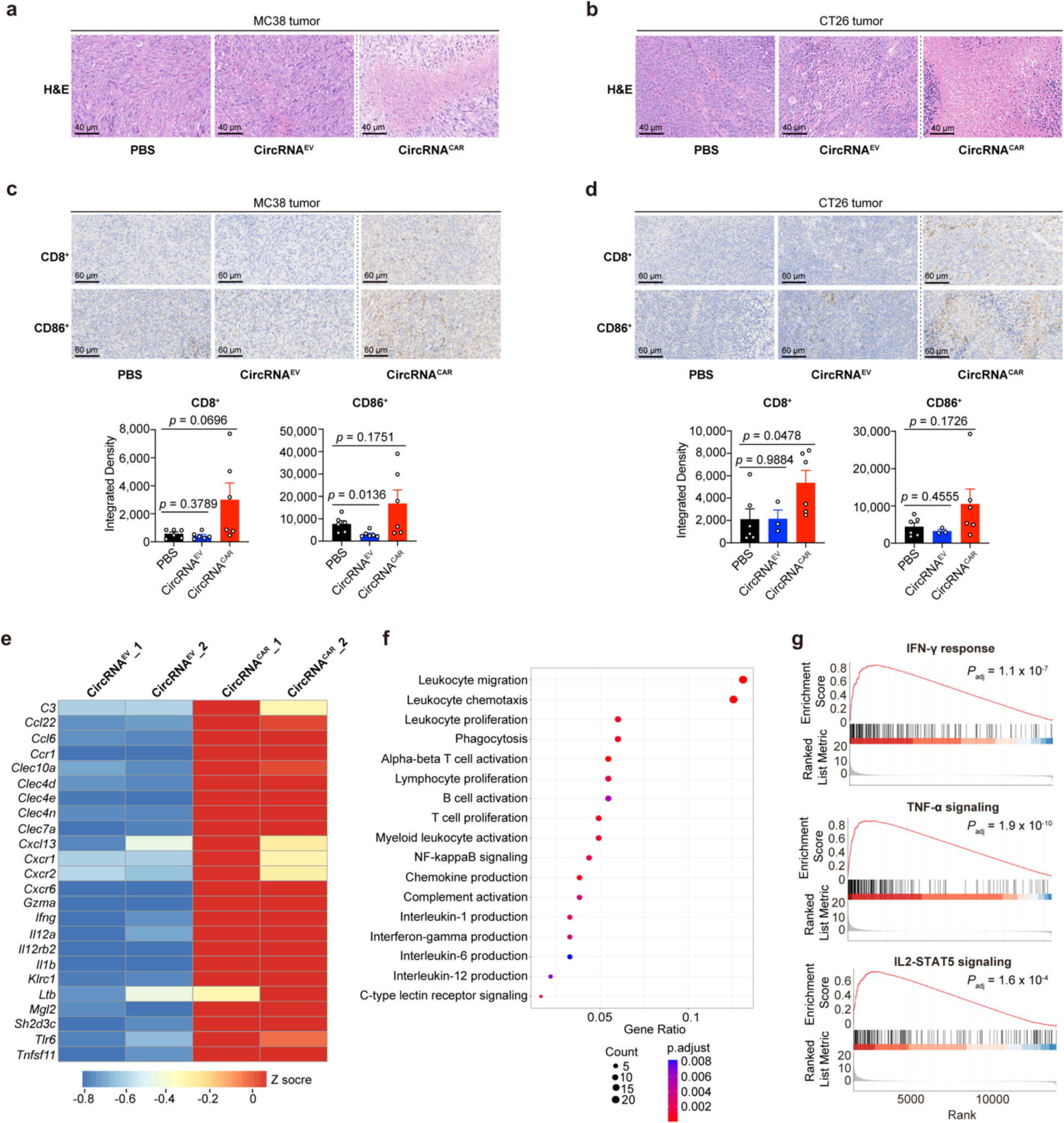
CircRNA^CAR^ reshaped the tumor microenvironment to a pro-inflammatory state in mice. **a**, **b**, H&E staining of tumor tissue sections obtained from MC38-HER2 (**a**) or CT26-HER2 (**b**) tumor-bearing mice after PBS, circRNA^EV^ or circRNA^CAR^ treatment via intratumoral administration. **c**, **d**, IHC staining of infiltrated CD8^+^ cells or CD86^+^ cells in tumor tissue sections obtained from MC38-HER2 (**c**) or CT26-HER2 (**d**) tumor-bearing mice after PBS, LNP-circRNA^EV^ or LNP-circRNA^CAR^ treatment via intratumoral administration. The integrated density of IHC staining quantified using ImageJ software. **e**, Heatmap of gene expression patterns of immune cells extracted from tumor tissues in the comparison between circRNA^EV^ and circRNA^CAR^ group via intratumoral administration (n = 2). **f**, Bubble chart of relevant biological processes involved in the circRNA^CAR^ group compared to the circRNA^EV^ group via intratumoral administration through GO analysis (n = 2). The size of the bubbles represented the number of genes, and the darkness indicated the significance. **g**, Gene Set Enrichment Analysis (GSEA) showing enriched pathways in the immune cells extracted from tumor tissues in the comparison between circRNA^EV^ and circRNA^CAR^ group via intratumoral administration (n = 2). In **c** and **d**, data were shown as the mean ± SEM, an unpaired two-sided Student’s t test was conducted for comparison, as indicated; *p < 0.05; **p < 0.01; ***p < 0.001; ****p < 0.0001; ns, not significant. Each symbol represents an individual mouse.

## Acknowledgements

We acknowledge other members of the Qu Laboratory for discussion throughout this study. We thank the Core Facility of Shanghai Medical College, Fudan University and the Public Technology Platform of School of Basic Medical Sciences, Fudan University for technical supports. This project was supported by funds from National Key R&D Program of China (2023YFC2604300 to L.Q.; 2022YFC3400100 to M.Q.); Key Project in Synthetic Biology of Science and Technology Commission of Shanghai Municipality (23HC1400100 to L.Q.); Morning Glory Plan of Shanghai Municipal Education Commission (to L.Q.); Research Project Plan of Shanghai Municipal Health Commission (20234Y0235 to L.Q.); National Natural Science Foundation of China (23DAA01060 to Y.X.); Shanghai Pujiang Program (23PJ1408900 to Y.X.).

## Author contributions

L.Q. conceived and supervised this project. L.Q., Y.W., L.L., X.W. and J. Li designed the experiments. Y.W., L.L., X.W., J. Li, F.G., J.Y., C. Zhang and Q.Y. performed the preparation of circRNAs and mRNAs, cell experiments, mouse experiments, detection experiments, and data collection with the help of J. Lin, D.L. and L.Q. C. Zhao, Y.X., X.W. and Y.W. performed the analysis of RNA-seq data. L.L., C. Zhang and Q.Y. prepared the LNPs with the help of M.Q. and L.Q. L.Q., Y.W., L.L., X.W. and J. Li wrote this manuscript with the help of other authors.

## Methods

### Plasmids construction

The anti-HER2/CD19 CAR and human HER2 antigen sequences were synthesized by Beijing Tsingke Biotech Co., Ltd. Subsequently, these sequences underwent double enzyme digestion and were cloned into the pUC57-circRNA-EV backbone using the Gibson assembly method, in preparation for subsequent in vitro transcription (IVT) reactions.

### Cell culture

HEK293T and CT26-WT cell lines were obtained from our laboratory, whereas THP-1, J774A.1, Jurkat, and SK-OV-3-Luc-EGFP cell lines were acquired from Shanghai Zhong Qiao Xin Zhou Biotechnology Co., Ltd. The HER2-overexpressing cancer cell lines, including MC38-HER2-Luc-EGFP, CT26-HER2-Luc-EGFP, 4T1-HER2-Luc-EGFP, and B16F10-HER2-Luc-EGFP were procured from Vigen Biotechnology (Zhenjiang) Co., Ltd. THP-1, Jurkat, SK-OV-3-Luc-EGFP, CT26-HER2-Luc-EGFP, 4T1-HER2-Luc-EGFP, B16F10-HER2-Luc-EGFP, and CT26-WT were cultured in RPMI 1640 medium, enriched with 10% fetal bovine serum and 1% penicillin-streptomycin. HEK293T, J774A.1, and MC38-HER2-Luc-EGFP cell lines were cultured in DMEM, supplemented with 10% fetal bovine serum and 1% penicillin-streptomycin. All cell lines were maintained in a 5% CO_2_ incubator at 37 °C.

### RNase R cleavage assay

The circRNA was heated at 65 ℃ for 3 min and cooled on ice. The RNase R (Epicenter, #RNR07250) was added and incubated at 37 ℃ for 5 or 15 min. 2 × RNA loading dye (NEB, #B0363S) were added to the system to stop the reaction and the RNA was separated by agarose gel electrophoresis.

### CircRNA transfection *in vitro*

Cells were seeded at a density of 3×10^5^ cells per well in 12-well plates for circRNA transfection. After 12 hours, cells were transfected with 2 µg of circRNA using Lipofectamine MessengerMax (Invitrogen, #LMRNA003) following the manufacturer’s protocol. Cells were harvested 24 hours post-transfection for further analysis.

### LNP encapsulation of circRNA

CircRNAs were encapsulated with lipid nanoparticles (LNP), referring to the methods reported in the previous literature^40^. CircRNAs were diluted with an acid buffer. The prepared LNP was mixed with circRNA solution at a volume ratio of 1:3. LNP-circRNA using NanoAssemblr Ignite Platform (Precision NanoSystems). And then the dialysis with 1×PBS buffer (pH = 7.2-7.4) and ultrafiltration was performed to concentrate LNP-circRNA with Amicon Ultracentrifuge Filtration device (Millipore). The concentration and encapsulation rate of circRNAs were measured using Quant-iT RiboGreen RNA Assay (Invitrogen, #R11490). The size and zeta potential of LNP-circRNA particles were measured by Zetasizer Pro (Malvern). For SORT lipid nanoparticles, the process of SORT-circRNA formulation referred to the method previously reported^23^. Briefly,the anionic lipids known as 18PA were dissolved in a solvent called tetrahydrofuran (THF). This solution was then blended with other additional lipids in ethanol, creating a mixture that contained the citrate buffer (10 mM, pH = 3.0). To prepare spleen-targeted SORT lipid nanoparticles containing 10% 18PA, a combination of lipids including 18PA, DOPE, Cholesterol, DMG-PEG2000, and C12-200, were prepared in ethanol to achieve a final molar ratio of 11.1:16:46.5:2.5:35^23^. This lipid blend was firstly mixed with 15 μL of ethanol, followed by the addition of 45 μL of circRNA solution containing 5 μg of circRNA, prepared in citrate buffer (10 mM, pH = 3.0). Blending the circRNA solution with lipid solution rapidly led to the formation of 10% 18PA SORT-LNP. Then the SORT-LNP formulayion undergone dialysis using SnakeSkin Dialysis Tubing (Invitrogen) for 2 hours and was then diluted with PBS for related experiments.

### Western blot

CAR expression in cells was assessed using Western blot analysis. Initially, cells transfected with circRNA for 24 hours were fractionated into supernatant and pellet. Pellet was resuspended with SDS loading buffer and heated at 100 °C for 10 minutes. Subsequently, proteins were resolved by SDS-PAGE and transferred onto PVDF membranes. The membranes were then blocked with 5% milk in PBST for 2 hours. The membranes were incubated overnight at 4 °C with detecting antibodies (Merck, #F1804). Afterward, the membranes were washed thrice with PBST, followed by incubation with HRP-conjugated goat anti-mouse IgG for 2 hours at room temperature. Finally, protein bands were visualized using ECL Western blot detection reagents.

### Flow cytometry analysis

For the detection of CAR expression, the expression of the chimeric antigen receptor (CAR) across different cells was evaluated via flow cytometry. Following 24 hours of RNA transfection, cells were centrifuged to collect the precipitates and washed thrice with PBS. Subsequently, cells were stained with a 1:50 dilution of PE-protein L (ACRO Biosystems, RPL-PP2H2). After a 30-minute incubation in the dark at 4 °C, the cells were washed with PBS three times. PE fluorescence was quantified using a Beckman Coulter CytoFLEX flow cytometer (AS03019).

For immunostaining analysis, firstly, the spleen or tumor tissues was isolated from mice, digested with collagenase solution at a final concentration of 1 mg/ml at 37°C for 30 minutes, then centrifuged to remove the supernatant, and washed twice with PBS before staining. For membrane surface staining, cells were stained with Live/Dead dye for 15 minutes, and then washed once with FACS buffer. Next, the staining buffer containing various membrane surface antibodies was added and the staining was kept in the darkroom at 4°C for 30 minutes. The intracellular staining was conducted according to the instruction of the eBioscience™ Foxp3/Transcription Factor Staining Kit. Briefly, after membrane surface staining, samples were washed once, and then fixed and permeabilized using a BD Cytoperm fixation/permeabilization solution kit according to the manufacturer’s instructions. Next, samples were washed once with FACS buffer, then washed with wash buffer, followed by intracellular staining in the darkroom at 4°C for one hour. After staining, samples were washed and acquired on Beckman Coulter CytoFLEX flow cytometer (AS03019). Analysis was performed using FlowJo software.

### *In vitro* killing assay

Seeded SK-OV-3-Luc-EGFP, MC38-HER2-Luc-EGFP, 4T1-HER2-Luc-EGFP, B16F10-HER2-Luc-EGFP and CT26-HER2-Luc-EGFP cell lines at a density of 2 × 10^4^ cells per well in a 96-well plate. Concurrently, co-cultured effector cells, transfected with circRNA, with the tumor cells in varying effector-to-target ratios. After 48 hours of co-culture, 100 µL of luciferase substrate was added to each well to assess the impact of THP-1, J774A.1, and Jurkat cells on the various tumor cell lines using a Microplate reader. In a similar manner, A549-EGFP-CD19 cells were seeded at a density of 2 × 10^4^ cells per well in a 96-well plate, and an equivalent volume of effector cells transfected with circRNA was added to the tumor cells, maintaining various effector-to-target ratios. After a 48-hour incubation, rinse the effector cells with Hank’s buffer, followed by the addition of 50 µL of XTT reagent. Incubate at 37 °C for 4 hours. Subsequently, using an ELISA reader to measure the absorbance at 450 nm with 650 nm as the reference wavelength.

### *In vitro* phagocytosis assay

Twelve hours post-circRNA transfection, different HER2^+^ tumor cells were co-cultured with THP-1 cells at various effector-to-target ratios for a 4-hour incubation. Following centrifugation at 1000 rpm, cells from each well were washed thrice with PBS and stained with APC/Cyanine7 anti-human CD14 Antibody (BioLegend, #123317) to assess THP-1 cell phagocytosis of HER2-positive tumor cells. After a 24-hour incubation with J774A.1 cells, cells were harvested, stained with either Brilliant Violet 510 anti-mouse/human CD11b Antibody (BioLegend, #101263) or PE/Cyanine7 anti-mouse/human CD11b Antibody (BioLegend, #101216), and incubated in the dark for 30 minutes at 4 °C, followed by three PBS washes. Subsequently, flow cytometry was employed to evaluate the phagocytic activity of effector cells against target cells.

### *In vitro* polarization assay

Forty-eight hours post transfection, the cells were harvested, washed with PBS, and subsequently stained with PE/Cyanine7 anti-mouse/human CD11b (BioLegend, #101216), PE anti-iNOS Antibody (BioLegend, #696806) and either Brilliant Violet 711 anti-mouse CD206 (MMR) Antibody (BioLegend, #141727) or Alexa Fluor 647 anti-human CD206 (MMR) Antibody (BioLegend, #321116) in dark. Following incubation, cells were pelleted by centrifugation at 1000 rpm. After triple PBS washes, the cells were incubated with antibodies in darkness at 4 °C for 30 minutes. Thereafter, following three additional PBS washes, macrophage polarization induced by circRNA was analyzed using flow cytometry.

### *In vivo* and *ex vivo* bioluminescence imaging

For intravenous administration, the female BALB/C mice (6-8 weeks) were injected intravenically with LNP-circRNA^luciferase^. 6 hours later, the luciferase substrate was intraperitoneally injected into mice, and then the luciferase imaging of the overall level or major organs including spleen, lung, liver, lymph node and kidney, was performed using the IVIS Lumina system (PerkinElmer, Lumina K). For intravenical administration, the female BALB/C mice (6-8 weeks) were injected subcutaneously with 1 × 10^6^ cells. When the tumor volume reached approximately 100 mm^3^, LNP-circRNA^luciferase^ or SORT-circRNA^luciferase^ was injected intratumorally. Then, the bioluminescence imaging was performed as described in the above intravenous administration.

### Vesicle detection

After circRNA^HER2-EPM-EABR^ transfection, cells were centrifuged for collecting supernatant at 24 h. Supernatant were filtrated by 0.45 μm filter and concentrated by 100 kD Amicon ultracentricentrifuge filtration device (Millipore). Vesicles were enriched by ultracentrifugation for 2 hours using Optima XPN-100 ultracentrifuge. The precipitation was adsorbed on the copper mesh after negative dyeing treatment and then detected vesicles by Thermo Scientific Talos L120C.

### Animal experiments

As for tumor-bearing mouse model, 6 × 10^5^ cancer cells, such as MC38-HER2 cells, CT26-HER2 cells or 4T1-HER2 cells, were subcutaneously inoculated into each C57BL/6 or BALB/c mouse.

When the tumor volume reached 80 mm^3^, mice carrying tumor were random grouped and received intratumoral injection with circRNA^CAR^ at a dose of 7.5 mg/kg every three days. Three mice in each group were euthanized after the last treatment, and the tumor tissues were sliced for H&E and IHC staining. And spleen and tumor tissues were used to analysis. To monitor the tumor volume, the tumor volume size was measured every other day. As for the combination therapy, when the tumor volume reaches approximately 80 mm^3^, they were randomly assigned to different groups for further analysis and injected circRNA^CAR^ (intratumorally or intravenously) plus circRNA^VAC^ (intramuscularly) at a dose of 7.5 mg/kg on day 10, 14, 22 after tumor engraftment. The mouse tumor volume was calculated using the formula: volume = (length × width^2^)/2.

### Animals and Ethics statement

The female BALB/c and C57BL/6 mice (6 to 8-week old) were ordered from Shanghai Model Organisms Center, Inc. All mice were bred and kept under specific pathogen-free (SPF) conditions in Shanghai Model Organisms Center, Inc. And all animal experiments were approved by the Committee on the Ethics of Animal Experiments of Fudan University and Shanghai Model Organisms Center, Inc., and undertaken in accordance with the National Institute of Health Guide for Care and Use of Laboratory Animals.

### Transcriptome-wide RNA-sequencing analysis

We extracted total RNA using the Quick-RNA^TM^ Miniprep Kit (ZYMO, # R1055), and the cDNA was generated using the NEBNext® Ultra™ II RNA Library Prep Kit for Illumina (New England Biolabs, #E7770S). Next, the DNA libraries were sequenced through the Illumina sequencing platform by Genedenovo Biotechnology Co., Ltd (Guangzhou, China). The raw FASTQ data underwent quality control analysis using FastQC^41^, with subsequent integration of results facilitated by MultiQC^42^. Trimmomatic^43^ was then applied to filter out low-quality reads and trim adapter content, employing parameters LEADING:3, TRAILING:3, SLIDINGWINDOW:4:20, and MINLEN:36. The filtered reads were aligned to the mouse reference genome (GRCm38) using HISAT2^44^, followed by sorting of alignment results and storage in BAM format using SAMtools^45^. Subsequently, featureCounts ^46^ was employed to generate the raw gene counts matrix, using parameters “-p --countReadPairs -t exon -g gene_id”. DESeq2^47^ was applied to identify differentially expressed genes, employing criteria of a p-value less than 0.01 and a logarithmic fold change greater than 2.5. Multiple testing correction was performed by Benjamini-Hochberg method. Gene set enrichment analysis (GSEA) was performed using clusterProfiler^48^, using HALLMARK and GO gene sets. Additionally, over-representation analysis (ORA) was conducted with GO and KEGG gene sets.

### Statistics

An unpaired two-sided Student’s *t* test was performed for comparison as indicated in the figure legends. Tumor growth curves were calculated by two-way ANOVA analysis as indicated in the figure legends. Survival curves were calculated by Kaplan-Meier simple survival analysis as indicated in the figure legends. *P < 0.05; **P < 0.01; ***P < 0.001; ****P < 0.0001; ns, not significant. Statistical analysis was performed with Prism 9 (GraphPad Software, Inc.).

## Data availability

All data and materials supporting the findings of this study in this manuscript are available from the corresponding author (L.Q.) upon a reasonable request under a completed Material Transfer Agreement.

